# spaDesign: A Statistical Framework to Improve the Design of Sequencing-based Spatial Transcriptomics Experiments

**DOI:** 10.1101/2024.09.10.612260

**Authors:** Juan Xie, Hyeongseon Jeon, Won Chang, Yeseul Jeon, Zihai Li, Qin Ma, Dongjun Chung

## Abstract

**Motivation:** Rigorous experimental design is essential for obtaining biologically meaningful findings and high-throughput spatial transcriptomics studies are not an exception. While some efforts have been made to improve the design of these studies, it is still significantly understudied yet how to optimize key experimental factors of these experiments. In the case of sequencing-based spatial transcriptomics studies, determining the minimum sequencing depth is an important experimental factor to decide.

**Results:** To address this critical limitation, here we propose spaDesign, a statistical framework to improve the design of sequencing-based spatial transcriptomics experiments. spaDesign is a statistically rigorously designed framework that employs Poisson Gaussian process and Fisher-Gaussian kernel mixture. It can easily simulate a range of spatial transcriptomics data with various sequencing depths, effect sizes, and spatial patterns, which allows rigorous estimation of needed total sequencing depths to detect spatial domains based on spatial transcriptomics experiments. We demonstrated the utility and power of spaDesign using 10X Visium data of the human brain and the chicken heart.

**Availability:** The R package and associated tutorial are freely available at https://github.com/JuanXie19/spaDesign.

## Introduction

High-throughput spatial transcriptomics (HST) technologies allow for the simultaneous measurement of gene expression and spatial locations of cells within tissues [Larsson et al., 2021, Rao et al., 2021]. The inclusion of spatial dimension provides an unprecedented opportunity to distinguish spatial domains, identify genes with varying spatial patterns, and make a more reliable prediction of cell-cell communications (CCCs). HST technologies have greatly enriched biomedical research and proven their clinical values [Yoosuf et al., 2020, Lyubetskaya et al., 2022, Chen et al., 2020].

The growing popularity of HST technologies generates a vast amount of HST data, presenting both opportunities and challenges in developing analytical methods and tools. Within the HST data analysis workflow, spatial domain identification is often the first step, as it forms the biological foundation for understanding spatial heterogeneity within complex organs [Liao et al., 2021]. Several computational methods and tools have been developed for spatial domain detection [Hu et al., 2021, Zhao et al., 2021] and comprehensively evaluated by Yuan et al. [2024]. Despite these advancements in analytical approaches, a notable gap remains regarding the design of HST experiments. One critical question in designing spatial transcriptomics experiments, particularly sequencing-based ones, is determining the required total sequencing depths to detect the target of interest. Although 10X Genomics has provided some recommendations [10x Genomics, 2024], they are not based on a strong statistical rationale. Recently, some efforts have been made to improve spatial transcriptomics experimental design by determining the physical locations on the tissue [Jones et al., 2024], and the number and size of the field of views (FoVs) [Bost et al., 2023, Baker et al., 2023]. However, these efforts have not specifically addressed the issue of selecting the optimal sequencing depth [Jeon et al., 2023].

On the other hand, existing tools for synthetic spatial transcriptomics data generation, such as SRTsim [Zhu et al., 2023] and scDesign3 [Song et al., 2024], may potentially be used for aiding experimental design by generating synthetic data. However, they do have some significant limitations for the aforementioned goals. Specifically, SRTsim preserves spatial gene expression patterns by maintaining the rank of a gene’s expression across spatial locations [Zhu et al., 2023]. Although this approach produces synthetic data that closely resembles reference data, its flexibility to customize the data according to users’ needs is limited, reducing its usefulness for experimental design purposes. scDesign3 models a gene’s marginal expected expressions as a smooth function of spatial locations and employs a Copula approach to handle correlations among genes [Song et al., 2024]. Although the simulated data closely mimics the reference data, its complex framework makes it difficult to generate data that deviates from the reference data in a controlled manner.

Hence, there is an urgent need to develop a robust statistical framework specifically tailored for HST experimental design. In response to this need, here we present spaDesign, a statistical framework for HST experimental design. SpaDesign integrates advanced statistical methods with a simulation-based framework, allowing easy generation of HST data with various sequencing depths, effect sizes, and spatial expression patterns. This enables a rigorous estimation of needed total sequencing depths for detecting spatial domains using HST experiments. Our framework equips researchers with the tools and guidance to conduct robust and reproducible HST experiments.

## Material and Methods

### SpaDesign overview

SpaDesign is a simulation-based statistical framework that aids researchers in designing HST experiments to achieve accurate spatial domain detection cost-effectively. The framework consists of four steps: feature selection, parameter estimation, data simulation, and performance evaluation. It expects users to provide pilot HST data with domain information. In Step 1, with that pilot data, spaDesign first selects domain-informative features. Then, in Step 2, it employs statistical models to estimate parameters that characterize the spatial patterns of the data. Given the parameters, in Step 3, spaDesign simulates gene expression data with different effect sizes and patterns. Finally, in Step 4, it applies spatial clustering tools to the simulated data to identify spatial domains and evaluate the performance. The workflow of spaDesign is shown in Fig 1, and details are provided in the following subsection.

**Fig. 1.**
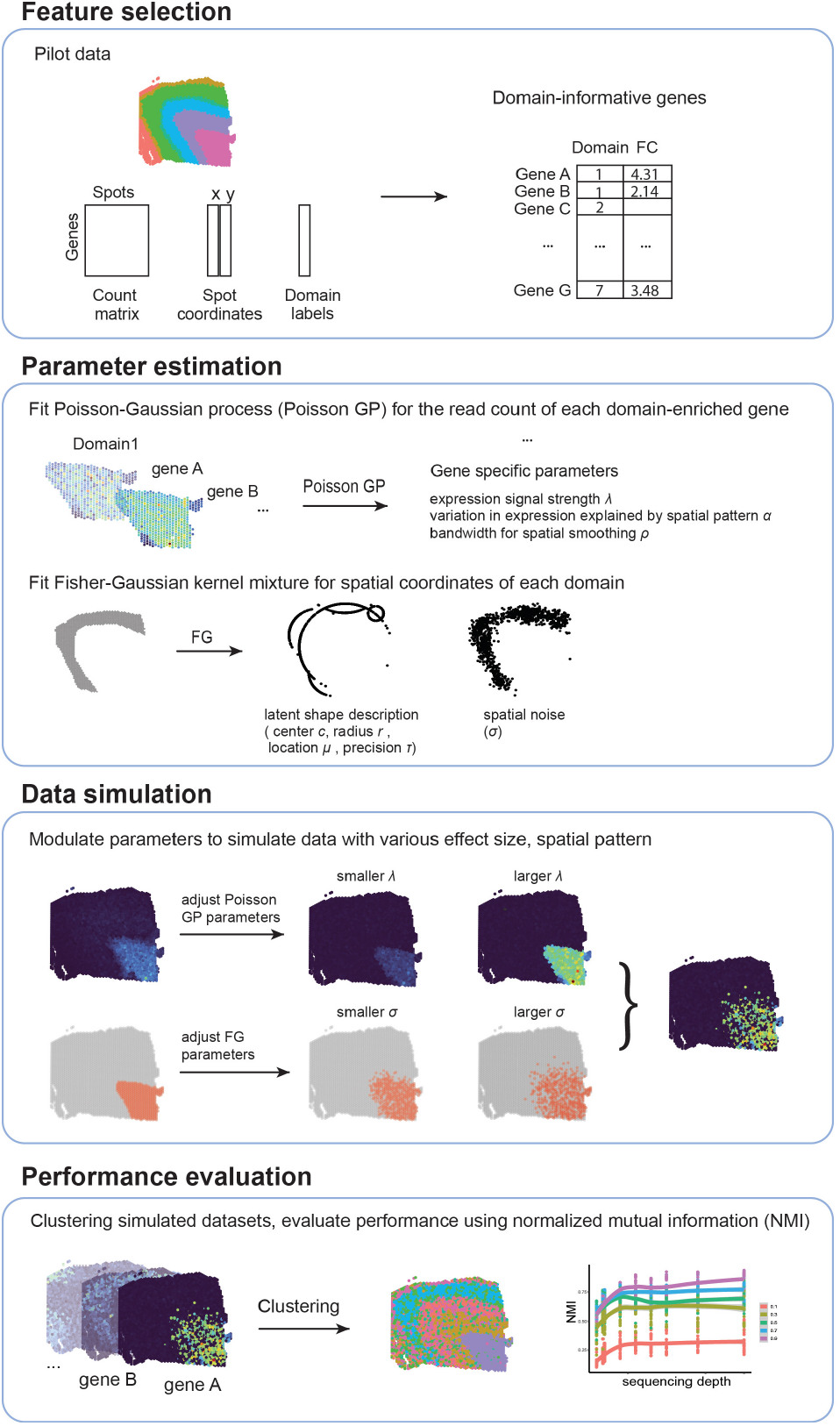
A schematic overview of spaDesign. spaDesign is a simulation-based framework for the design of spatial transcriptomics experiments, which consists of four steps: feature selection, parameter estimation, data simulation, and performance evaluation.

#### Step 1: Feature section

For a given pilot dataset, genes with high absolute fold change are selected as domain-informative features. Specifically, consider a tissue with multiple domains, each containing a certain number of cell spots. The absolute fold change for a gene is calculated by comparing its average expression within a domain to its expression in the lower 50% of spots outside that domain. Note that when selecting the spots outside the domain, we consider only the lower 50% of spots to avoid the impact of spots with ambiguous expressions. SpaDesign selects genes that exceed a user-specified threshold for absolute fold change, identifying those that are highly informative for a specific domain.

#### Step 2: Parameter estimation

In step 2, the Poisson Gaussian process (Poisson GP) and Fisher-Gaussian (FG) kernel mixture models are used to estimate parameters that characterize the spatial pattern in gene expression and domain location distribution, respectively.

### Poisson Gaussian process model

Poisson GP is a spatial generalized linear mixed model commonly used to model spatially varied count data [Diggle et al., 1998, Wakefield, 2007]. In our framework, we utilize this model to describe the gene spatial expression patterns. Specifically, for a given gene *g* and tissue domain *d* with *n*_*d*_ spots, let ***S***_***i***_ = (*s*_*i*1_, *s*_*i*2_)’, *i* = 1, …, *n*_*d*_, denote the normalized spatial coordinates for cell spot *i*, and *y*_*i*_ (*s*_*i*_) denote the gene expression (read counts) for location *i*.

We define a Poisson GP as follows:

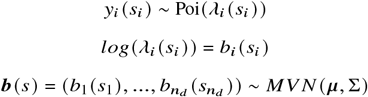

Here *λ*_*i*_ (*s*_*i*_) is an unknown rate parameter denoting the underlying gene expression level for gene *g* at spot *i*. The log scale of *λ*_*i*_ (*s*_*i*_), *b*_*i*_ (*s*_*i*_), is assumed to follow a stationary Gaussian process model with mean *μ* and covariance Σ. The spatial dependency among the gene expression across spatial locations is captured by *b*_*i*_ (*s*_*i*_), or more precisely, the covariance function Σ. In SpaDesign, we adopt the absolute exponential kernel for Σ, i.e., for two spots *s*_*i*_ and *s* _*j*_,

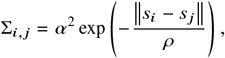

where 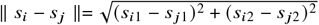 is the the Euclidean distance between *s*_*i*_ and *s* _*j*_. *ρ* controls how fast the correlation in the expression between two locations decays, and the larger *ρ*, the stronger the correlation. *α* describes the portion of gene expression variation that can be attributed to spatial locations. A Bayesian framework with Markov Chain Monte Carlo (MCMC) technique is used to estimate the parameters. For mean *μ*, we opt for weakly-informative priors by assuming *μ* ∼ *N* (0, *sd* = 100). For *ρ*, we assume *ρ* ∼ inverseGamma(*α*_*ρ*_, *β*_*ρ*_), where *α*_*ρ*_, *β*_*ρ*_ are the shape and scale parameters, respectively. Here we choose *α*_*ρ*_ = *β*_*ρ*_ = 5. This prior distribution ensures that the support of *ρ* to be positive and discourages its value to be too close to zero. For the scale factor *α*, we adopt a stronger prior by assuming *α* ∼ *N* (0, 0.1) to address potential identifiability issues.

### Fisher-Gaussian kernel mixture model

FG kernel mixture model was initially proposed for multivariate density estimation in non-linear manifolds [Mukhopadhyay et al., 2020]. The core of this mixture model is the FG kernel, which adds Gaussian noise to von Mises-Fisher density [Fisher et al., 1993] to model Gaussian density concentrated near the circumference of a sphere [Mukhopadhyay et al., 2020]. By plugging the FG kernel in mixture models, it provides a flexible, parsimonious, and accurate density approximation. In our framework, we use this model to characterize the spatial point pattern of the tissue domains by describing latent spatial patterns via a collection of partial circles with different centers and sizes. The details are as follows.

For a given tissue domain *d* with *n*_*d*_ spots, let ***s***_***i***_ = (*s*_*i*1_, *s*_*i*2_)’, *i* = 1, …, *n*_*d*_, denote the normalized spatial coordinates for cell spot *i*. We represent the distribution of ***s*** using a FG kernel mixture model:

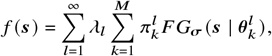

where 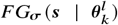 is the FG kernel parameterized by 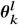, and 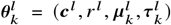 is a set of latent spatial shape parameters. ***c***^*l*^ and *r* ^*l*^ denote the center and radius of the sphere, 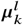 is the direction in which the density is most concentrated, 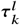 controls the degree of concentration, and ***σ*** controls the Gaussian noise level. *λ*_*l*_ is the second-layer mixing weight and 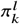 is the mixing weight for the *k*-th FG kernel within *l*-th layer. The FG kernel takes the following form (we drop the superscripts and subscripts for simplicity) :

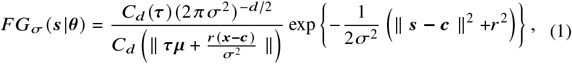

where 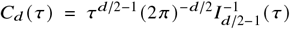, and *I*_*ν*_ denotes the modified Bessel function of the first kind of order *ν*.

For parameter estimation, a Bayesian framework using MCMC sampling proposed by Mukhopadhyay et al. [2020] is employed. In short, it uses a Dirichlet process prior for the mixing weights ***λ*** = (*λ*_1_, *λ*_2_, …,), and a Dirichlet prior for the FG kernel weights 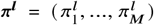. Conjugate priors are assigned to the parameters of the von Mises-Fisher parameters 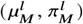 with 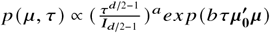. An inverse gamma prior is assigned to the Gaussian noise *σ*^2^. For the details of inference procedure, please refer to Mukhopadhyay et al. [2020].

#### Step 3: Data simulation

Based on the estimated parameters obtained from Step 2, spaDesign simulates read counts in a domain-specific manner. For a given domain *d* with *n*_*d*_ spots, the simulation begins with domain-informative genes. Specifically, for each domain-informative gene, spaDesign first simulates the read counts across these *n*_*d*_ locations (within-domain counts) based on the fitted Poisson GP. Next, the read counts outside the domain are simulated using a Poisson model, where the mean of the lower 50% of outside-domain spots is used as the parameter of the Poisson model. This process is repeated for all domain-informative genes, resulting in a complete set of read counts for domain *d*. By applying this procedure across all domains, spaDesign generates the full read count profile for the entire tissue.

When SpaDesign simulates within-domain read counts, spaDesign employs a conditional sampling approach. Specifically, for a given tissue domain *d* with *n*_*d*_ spots, let ***s***_***i***_ = (*s*_*i*1_, *s*_*i*2_)’ denote the normalized spatial coordinate for spot *i*, and *y*_*i*_ (*s*_*i*_) denote the gene expression count for gene *g* across the spots (we drop the subscript for *g* for simplicity). To balance resemblance and variation, we randomly select 70% spots among these *n*_*d*_ spots, and do conditional sampling for the *n*_*d*_ spots to simulate *y*_*i*_ (*s*_*i*_). To distinguish, we let ***S*** to denote the original spots, and use *D*_*k*_ = (*S*_*k*_, *b* (*S*_*k*_)) to denote the set of selected spots (both location and gene expression values). Then, for all spots in ***S***, we jointly simulate their read counts, conditional on the read counts at the spots in *D*_*k*_ :

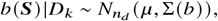

where 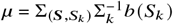 and 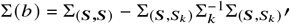.

After obtaining *b*_*i*_ (*s*_*i*_), we further generate *y*_*i*_ (*s*_*i*_) based on Poisson model with *λ*_*i*_ (*s*_*i*_) = *exp* (*b*_*i*_ (*s*_*i*_)). This conditional sampling approach ensures that the simulated within-domain read counts capture the spatial dependencies within domain *d*. Note that when we assessed the impact of the number of selected spots (|*S*_*k*_ |) on the read counts simulated using the conditional sampling, we found that the simulated expression generally resembles the original expression when |*S*_*k*_ | is between 50% and 100% of *n*_*d*_, and it starts to significantly deviate from the original expression when |*S*_*k*_ | is below 50%. Based on this rationale, we decided to use an intermediate proportion, i.e., selecting 70% of the spots for conditional sampling. For details, please refer to the Supplementary Materials (Section 1, Fig S1).

In addition, spaDesign also allows users to simulate data with modified spatial expression patterns. To accomplish this, spaDesign utilizes the fitted FG kernel mixture model. By controlling the Gaussian noise parameter *σ*^2^, we can adjust the dispersion of the generated spatial locations, thus influencing the spatial expression pattern of genes. After generating within-domain read counts for the original *n*_*d*_ locations, spaDesign creates a set of *n*_*d*_ new locations based on the fitted FG kernel mixture model. Subsequently, it establishes a one-to-one correspondence between the set of new locations and the original locations using the Hungarian algorithm [Kuhn, 1955]. Finally, the read counts associated with the original locations are transferred to the new locations.

During data simulation, we can easily modify the sequencing depth, gene expression level, and spatial expression patterns by adjusting the estimated parameters, which facilitates the evaluation of spatial domain prediction performance under different scenarios. The simulated data inherits the domain labels from the pilot data.

#### Step 4: Performance evaluation

After the data simulation, we predict spatial domains by applying SpaGCN [Hu et al., 2021] to the simulated data. To quantify the similarity between the inferred domain labels and the ground-truth domain labels, we compute the normalized mutual information (NMI) metric [Strehl and Ghosh, 2002], which has been used to evaluate the performance of spatial domain identification tools [Yuan et al., 2024]. NMI ranges from 0 to 1, with a higher value indicating better agreement between the clustering labels. While the Adjusted Rand Index (ARI) is also commonly used to assess clustering results, it focuses more on the agreement or disagreement of cluster assignments rather than the overall structure of the clusters. In contrast, NMI considers both the cluster assignments and the distribution of samples within clusters, making it generally more robust to imbalances in cluster sizes, compared to ARI. Therefore, we adopt NMI for our evaluation. Note that in addition to SpaGCN, we also checked the results using two other spatial domain prediction tools, BayesSpace [Zhao et al., 2021] and Louvain [Blondel et al., 2008]. The details of these methods and the corresponding results can be found in the Supplementary Materials (Sections 2 and 5, Fig S5 and S6).

### Pilot datasets

To illustrate the proposed statistical framework, in this paper, we investigated two HST datasets with manual domain annotations as pilot data. Both datasets were generated using the 10x Genomics Visium platform. The first dataset is from the human dorsolateral prefrontal cortex (DLPFC), obtained from spatialLIBD [Pardo et al., 2022], which is widely used to benchmark spatial clustering tools [Shang and Zhou, 2022, Yuan et al., 2024]. This dataset contains 12 samples from three adult donors. In our analysis, we focused on sample 151673, which contains expression profiles of 33,538 genes across 3,639 spots. According to the original study [Maynard et al., 2021], most locations were clustered into one of seven tissue domains. Here we focused on locations with domain annotation information (3,611 spots) and discarded the remaining unannotated spots. The second dataset is from chicken heart tissue, sourced from Mantri et al. [2021]. This dataset comprises 12 heart tissue sections collected at four key development stages of chicken hearts. Here we focused on the sample collected on day 14, which contains measurements of 24,356 genes across 1,967 spots on the tissue. The study originally clustered these locations into one of six anatomical regions. In addition, we also investigated two additional human brain datasets, the details and results of which can be found in the Supplementary Materials (Section 4, Fig S3 and S4).

### Simulation scenarios

#### Simulate data with different effect sizes

To examine the effects of effect size on spatial domain detection performance under different sequencing depths, we explored six choices of total sequencing depths (50%, 100%, 300%, 500%, 700%, and 1,000% of the observed total sequencing depth), and then for each sequencing depth, we further explored five choices of effect size (0.5, 1, 3, 5, and 7 times of the original effect size). The above adjustment was achieved by modifying the *λ*_*i*_ in the Poisson GP. For each sequencing depth and effect size combination, 100 simulation replicates were performed.

#### Simulate data with different spatial expression patterns

To investigate the effects of spatial expression patterns on spatial domain detection performance, we simulate datasets with different degrees of dispersion. To do so, we randomly selected a proportion of genes from the set of domain-specific genes, and disturbed their spatial expression pattern by using larger *σ* values of the FG kernel mixture models during data simulation (1 and 3 times the estimated *σ*). For the remaining genes, we kept their spatial expression pattern unchanged. We varied the proportion of genes with disturbed spatial expression patterns (from 10% to 90%). To avoid the bias caused by the randomness during gene selection, we repeated the simulation 1,000 times for each proportion and used the average of the resulting 1,000 count matrices for spatial domain detection. Besides, to examine the interaction between sequencing depth and spatial patterns, we introduced different sequencing depths during simulation (50%, 90%, 100%, 300%, 500%, 700%, and 1,000% of the original total sequencing depth). For each proportion and sequencing depth combination, 100 simulation replicates were performed.

## Results

### Ability of spaDesign to resemble and modify real spatial transcriptomics data

The parameter estimation and the data simulation serve pivotal roles in our simulation-based statistical framework. To provide realistic and reliable guidance for experimental design, a good framework should be able to simulate data that mimic the characteristics of real data. To examine how well spaDesign can capture spatial patterns in HST data, we investigated two real HST data produced by the 10x Visium platform, including the human dorsolateral prefrontal cortex (DLPFC) data (referred as human brain data thereafter) [Maynard et al., 2021] and the chicken heart data collected on the late four-chamber development stage (referred as chicken heart data thereafter) [Mantri et al., 2021]. Since the results are generally consistent across the two datasets, here we showcase the results for the human brain data, while the results for the chicken heart data are provided in the Supplementary Materials (Fig S2).

Fig 2 demonstrates how well spaDesign can retain spatial patterns using four domain-specific genes. Fig 2A shows that both the overall gene spatial expression patterns and the magnitude of the expression values in the pilot data are well reserved in the simulated data. This phenomenon holds regardless of the shape and range of the spatial pattern. Fig 2B further indicates that spaDesign effectively models the diverse spot distribution patterns and accurately simulates spots with similar distribution patterns as those observed in the pilot data.

**Fig. 2.**
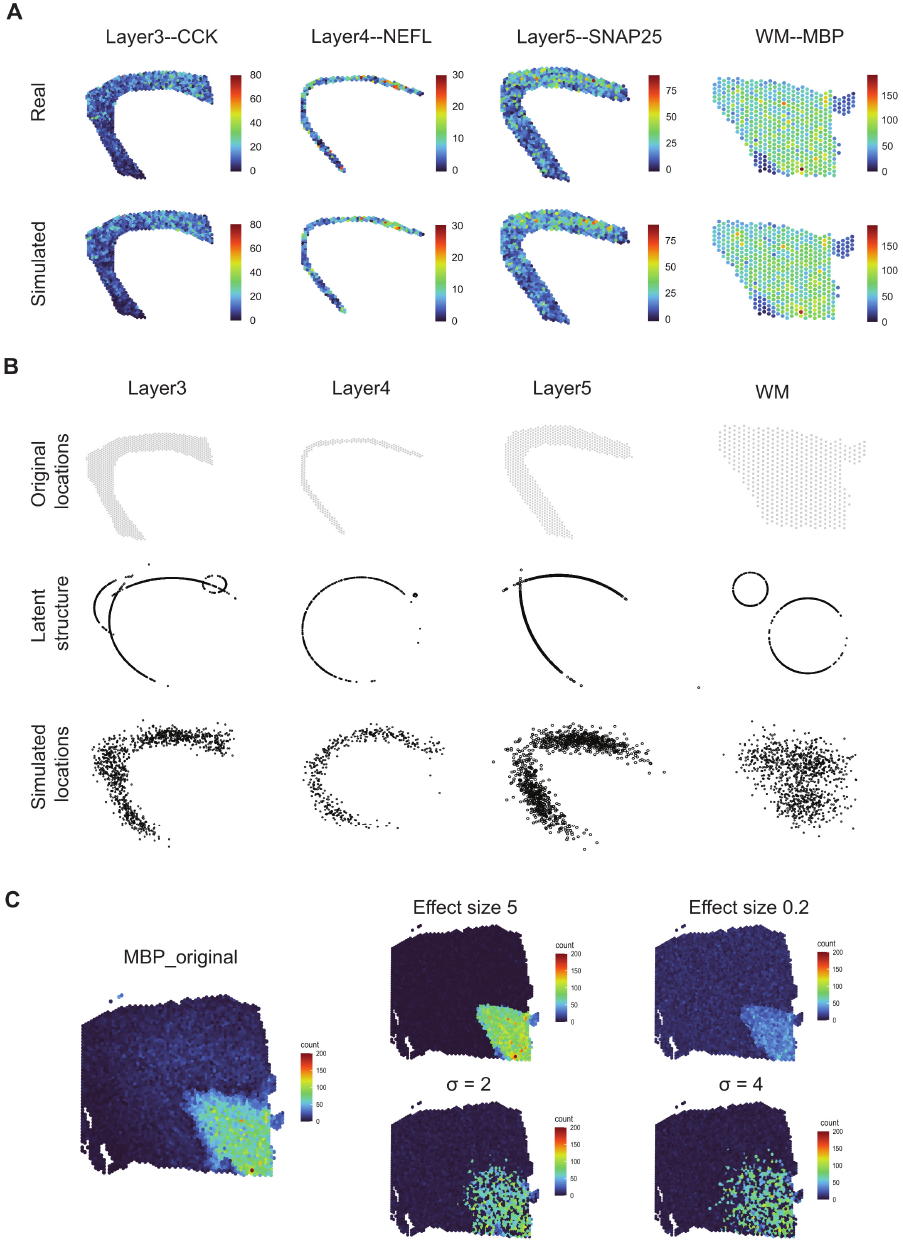
spaDesign has powerful capability to simulate spatial transcriptomics data in a flexible way. (A) The spatial expression patterns of four representative genes are displayed for the pilot data (the first row) and the simulated data generated by spaDesign (the second row). (B) The spatial distribution of points from four selected domains is displayed (the first row), along with the estimated latent structure (the second row) and the simulated locations generated using spaDesign (the third row). (C) The overall expression for MBP in the pilot data is displayed (left), along with the simulated MBP expression with stronger/weaker expression signal (first row on the right), and those with dispersed spatial pattern (second row on the right).

In addition to simulating datasets that mimic the pilot data, another crucial feature of spaDesign is the ability to introduce controlled levels of noise or variability into the simulated data. This capability is illustrated in Fig 2C. Specifically, by modifying a single parameter in the Poisson GP model, we can obtain a dataset with either a stronger or weaker signal compared to the pilot data. Likewise, by adjusting a single parameter in the FG kernel mixture model, we can easily generate a dataset with a more dispersed spatial expression pattern.

This feature enables method developers to thoroughly assess the robustness and generalizability of their analytical methods under various conditions and scenarios. This feature is also important from an experimental design perspective. Real-world data are often noisy. Simulating data with controlled noise levels enables researchers to evaluate the sensitivity of analytical methods to detect true signals amidst varying degrees of noise. By assessing the impact of noise on the performance of analytical methods, researchers can make informed decisions about experimental design to maximize the efficiency and effectiveness of their studies. In the end, it allows researchers to determine the minimum requirements needed to achieve reliable results.

spaDesign’s flexible ability to simulate data with similar or noisier spatial patterns than the pilot data, is attributed to the two embedded statistical models. In particular, the Poisson GP mainly deals with the spatial expression pattern, while the FG kernel mixture model handles the distribution of spot locations. The parameters in the two models have clear interpretations, allowing easy data generation with various characteristics. We note that such capability to introduce controlled noise is missing in existing tools that simulate HST data. Hence, we believe that spaDesign has its own unique contribution to the HST data simulation and the design of HST experiments.

### Impact of effect size on the spatial domain detection performance

Regional heterogeneity of gene signatures is widely acknowledged and can change during aging, infection, and many other biological processes [Tan et al., 2020]. For instance, it is reported that aged mice had greater upregulation of microglial activation markers (e.g., CD11b, CD68) in the white matter compared with the gray matter [Hart et al., 2012]. This regional heterogeneity, described by effect size, is an important biological factor affecting tissue architecture detection performance. To understand how the effect size can affect the spatial domain detection performance, we simulated data with various effect sizes under multiple total sequencing depths using the human brain data and the chicken heart data as pilot data (Fig 3A). Then we identified spatial domains from the simulated datasets using SpaGCN, and evaluated the performance using the NMI metric. For each simulation scenario, 100 replicates were generated. The LOESS smoothing method [Cleveland and Devlin, 1988] was applied to describe overall relationships between the detection performance and the total sequencing depth.

**Fig. 3.**
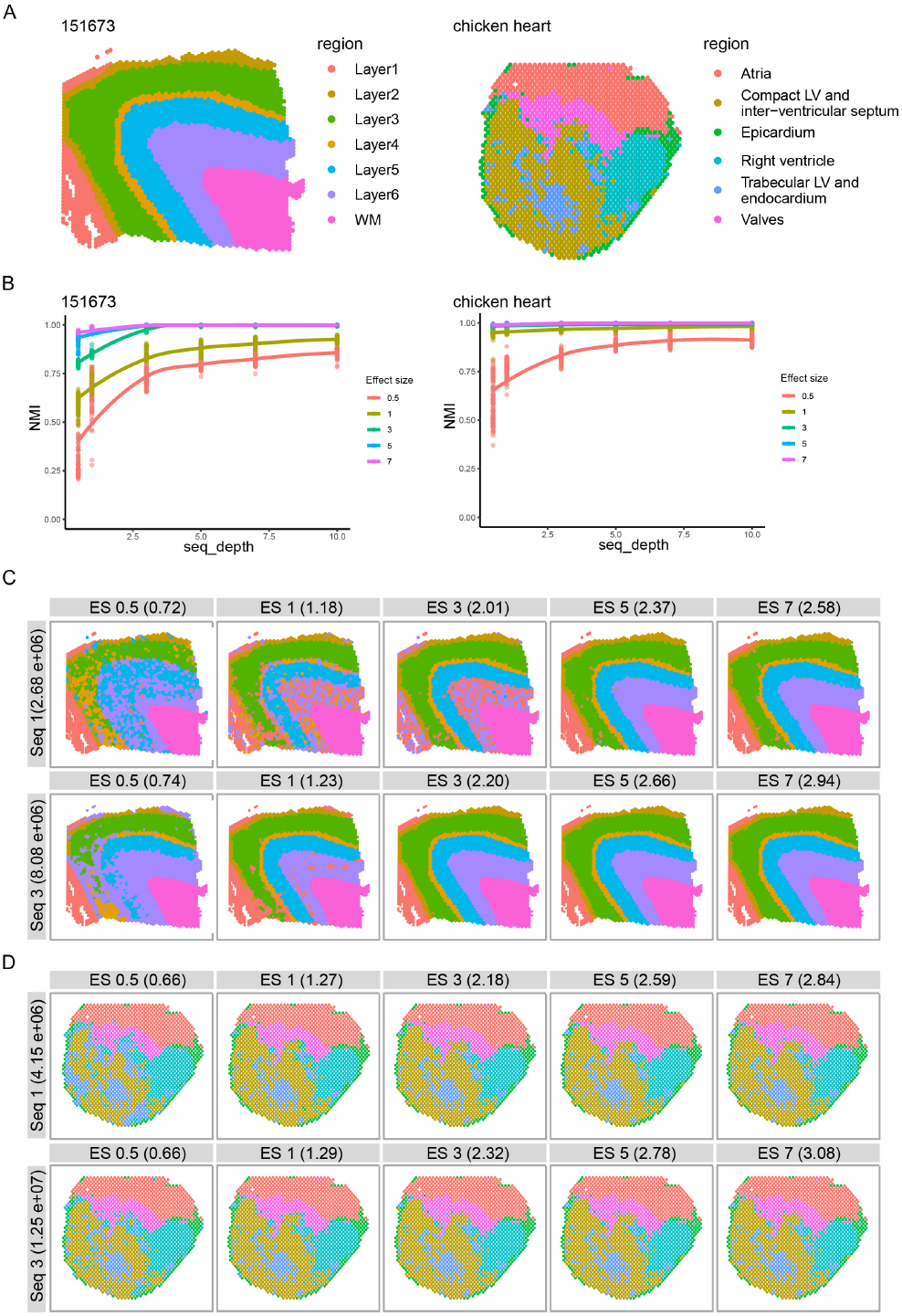
Impacts of effect sizes on the spatial domain detection performance. (A) Manual annotations for the two real datasets. The sample 151673 from the human brain DLPFC data contains seven layers, including six DLPFC layers and white matter (WM). The chicken heart D14 data contains six anatomical regions. (B) NMI curves as a function of sequencing depths (*x* axis) under different effect sizes (lines) for the human brain data (left) and the chicken heart data (right), respectively. (C) and (D) show the example spatial domain detection results produced by SpaGCN on the simulated data under different effect sizes (columns) and sequencing depths (rows) for the human brain data (C) and the chicken heart data (D), respectively. The numbers in the parenthesis denote the averaged absolute fold change.

Consistent with expectations, we observe that larger effect sizes lead to higher NMI, and larger sequencing depths also give rise to higher NMI, for both the human brain data and the chicken heart data (Fig 3B). Specifically, for smaller effect sizes, as the sequencing depth increases, NMI increases rapidly at the beginning, and then it reaches a plateau. For larger effect sizes, the curves look like a flat line, i.e., we can achieve the potentially maximal NMI with small sequencing depths. In other words, these results indicate that the spatial domain detection performance would finally saturate as sequencing depth increases, and the larger the effect size, the earlier the saturation. These observations make natural sense because increasing the sequencing depth can potentially improve signal-to-noise ratios (SNR). On the other hand, when the effect size is large enough, the potential maximal SNR is already achieved with the small sequencing depth, thus further increasing the sequencing depth would not provide additional benefit.

To further confirm this explanation, we showcase some example spatial domain detection results under different combinations of sequencing depths and effect sizes (Fig 3C and 3D). Here we can observe that in the case of low effect sizes (those on the left side in Fig 3C and 3D), although the major regions in the spatial domains were captured, still a significant number of spots were misclassified. As effect size increases, the number of misclassified spots decreases significantly (those on the right side in Fig 3C and 3D). In addition, compared to the lower sequencing depth (the first row in Fig 3C and 3D), we could observe a significantly less number of misclassified spots with the higher sequencing depth (the second row in Fig 3C and 3D). We believe that these examples confirm our explanation of the observations for Fig 3B.

During this investigation, we recognized interesting differences between the human brain data and the chicken heart data, although the overall relationships between NMI, sequencing depth, and effect size remain similar. Specifically, in the case of human brain data, the effect size comparable to three times that of the pilot data was needed to make NMI reach early saturation. In contrast, for the chicken heart data, the effect size equivalent to that of the pilot data already made NMI reach early saturation. Several factors can contribute to this difference. Firstly, the SNR of the chicken heart data is generally higher than that of the human brain data (absolute fold change: 1.27 vs. 1.18), which implies differences in the baseline of SNR between the two datasets. Secondly, two datasets are also different in the sense of the spatial structure complexity. Specifically, the human brain data exhibits a layered spatial structure with varying layer widths, whereas the chicken heart data comprises three large chunks and three smaller ‘patches’. These might indicate that spatial domain detection performance can be affected by various characteristics of the tissue under consideration. In practice, this implies the importance of selecting appropriate pilot data when designing experiments. In this sense, spaDesign can support more accurate experimental design by using the pilot data that are more similar to the tissue that will be used in the to-be-implemented experiments.

We note that the NMI curves obtained using spaDesign also have important implications for the design of HST experiments. Here, the total sequencing depths of the human brain data and the chicken heart data were approximately 16 million and 19 million reads, respectively. According to the guidelines provided by 10X Genomics, the minimum total sequencing depth for these samples should be 90 million and 49 million reads, respectively. However, according to Fig 3B, when the effect size is comparable to that of the human brain data, the curve made a plateau at around 3 times the original sequencing depth, which is approximately 48 million reads. In the case of the chicken heart data, when the effect size is comparable to that of the chicken heart data, the curve already made a plateau at the original sequencing depth, equivalent to 19 million reads. For both the human brain data and the chicken heart data, we found that spaDesign suggests a total sequencing depth that is significantly lower than what the 10X recommended. This might imply that the guidelines provided by 10X Genomics may overestimate the required sequencing depth for these two datasets. Hence, we believe that spaDesign can help researchers for their HST experimental designs by making a better balance between the data quality and the cost-effectiveness.

### Impacts of spatial expression pattern on the spatial domain detection performance

Gene spatial expression pattern is another important biological factor that can affect spatial domain detection performance. To examine how spatial domain detection can be affected by gene spatial expression patterns, we introduced various degrees of spatial noise to the simulated data, to disturb the gene’s spatial expression pattern. Specifically, for each spatial noise level, we varied the proportion of genes with disturbed spatial patterns, as well as the total sequencing depth, and applied SpaGCN to the simulated datasets.

For the human brain data, we observed a consistent trend in the NMI across sequencing depths, irrespective of the proportion of disturbed genes (Fig 4A): the NMI initially increases with sequencing depth before reaching a plateau. Additionally, we discern a distinct segregation of the NMI curves obtained under varying proportions of disturbed genes. Specifically, the NMI curve obtained at a high proportion of disturbed genes is notably lower compared to those obtained at lower proportions. As the proportion of disturbed genes decreases, the NMI curves converge. Similar trends were also observed for the chicken heart data (Fig 4A). Fig 4B and 4C showcase several spatial domain detection results under different combinations of total sequencing depths and proportion of disturbed genes for the human brain data and the chicken heart data, respectively. When there is a higher proportion of disturbed genes (those on the left side in Fig 4B and 4C), the predicted spatial domain looks more chaotic, i.e., it is harder to achieve clear spatial separation. As the proportion of disturbed genes decreases (those on the right side in Fig 4B and 4C), spatial domains can be better distinguished. When looking at the results of the same proportion of disturbed genes but from different total sequencing depths (the first vs. the second rows in each of Fig 4B and 4C), we could see that increasing the total sequencing depth improves the performance to some degree.

**Fig. 4.**
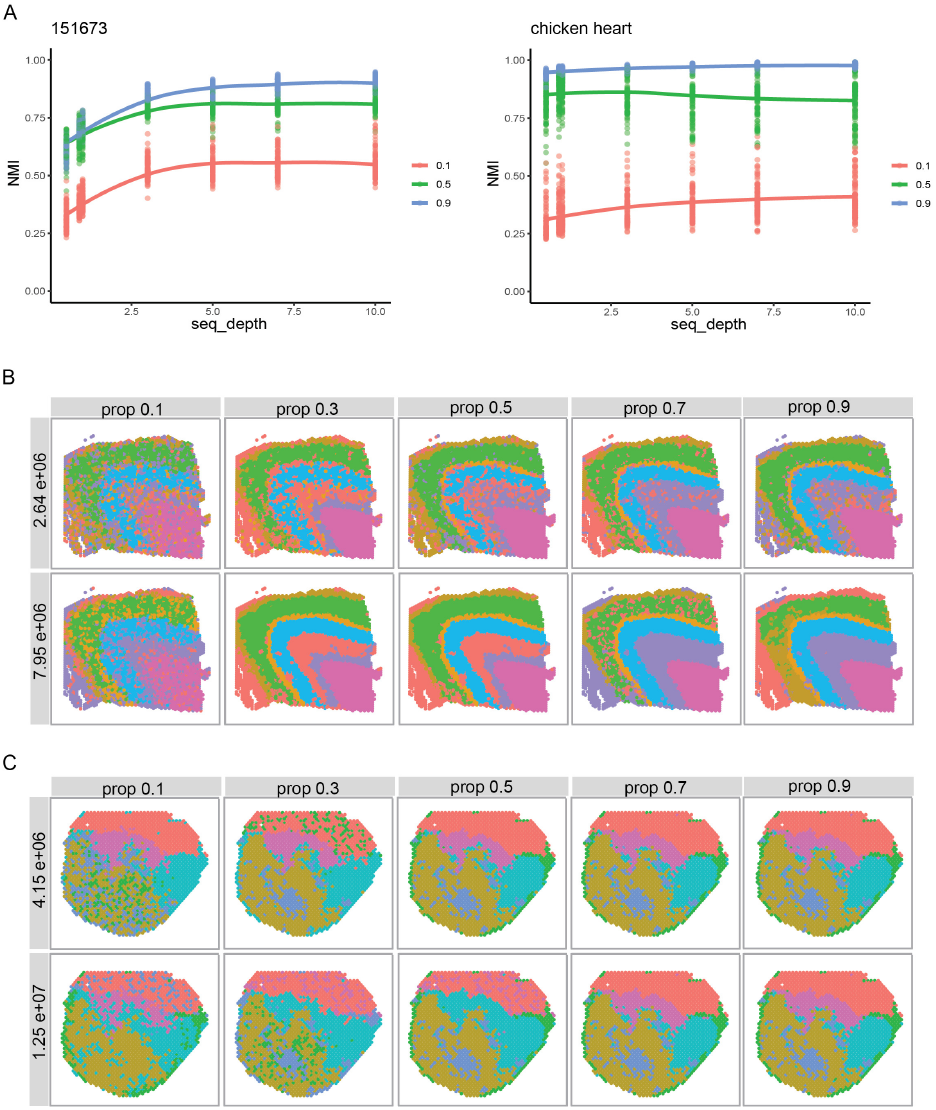
Impacts of spatial expression patterns on the spatial domain detection performance. (A) NMI curves as a function of sequencing depths (*x* axis) for different proportions of disturbed genes (lines) for the human brain data and the chicken heart data, respectively. (B) and (C) show the example spatial domain detection results produced by SpaGCN on the simulated data under different proportions of disturbed genes (columns) and sequencing depths (rows) for the human brain data (B) and the chicken heart data (C), respectively.

These results underscore the impact of gene spatial expression patterns on the detection of spatial domain detection. To improve the spatial domain detection, many tools (including SpaGCN) leverage spatial proximity information among spots. As a result, if the gene spatial expression pattern aligns with the underlying spatial domain, leveraging spatial proximity information will improve the spatial domain detection. For the same reason, disruption of spatial expression patterns, which leads to divergence from the underlying spatial domain structure, can degrade the spatial domain detection performance. The results here may carry practical implications. For instance, tumors often exhibit greater heterogeneity in gene expression due to factors such as the presence of various cell types within the tumor microenvironment. This increased ‘noise’ in gene expression patterns can pose greater challenges for identifying coherent spatial structures within the tumor tissue compared to normal tissues. Consequently, when working with tumor samples, one may anticipate a likely poorer performance in spatial domain detection compared to healthy samples, even under similar total sequencing depth and tissue type. Furthermore, while increasing the total sequencing depth may improve the detection, it is possible that it cannot fully remedy this issue.

## Discussion

In this paper, we presented spaDesign, a simulation-based statistical framework tailored to assist in the design of HST experiments. spaDesign offers a statistically rigorous and flexible approach to simulate data closely resembling the pilot data. It does not only allow researchers to investigate the impacts of sequencing depths and effect sizes, but also the impacts of gene spatial expression patterns. These features allow a systematic evaluation of the impact of sequencing depth, effect size, and spatial expression pattern on the spatial domain detection performance. Notably, our framework provides insights on the minimum required total sequencing depth for detecting spatial domains for a given tissue. Take the human brain data for example. While 10X Genomics suggested a minimum of 25,000 read pairs per spot covered with tissue [10x Genomics, 2024], equivalent to the sequencing depth of 90 million reads, spaDesign suggests the sequencing depth of 48 million reads, which is much lower than what is recommended by the 10X Genomics. Hence, we believe that spaDesign can potentially be a powerful tool that can better support researchers in their design of HST experiments.

Looking ahead, we plan to improve spaDesign from various perspectives. First, we are exploring ways to expedite the implementation of the framework. Currently, both the Poisson GP and the FG kernel mixture model employ the MCMC for parameter estimation, which may take a long time to run. In addition, the computation time to fit a Poisson GP also increases as the number of spots increases. To address this, we are currently investigating various alternatives such as the nearest-neighbor Gaussian process [Datta et al., 2016, Liu et al., 2020], variational inference [Blei et al., 2017], and approximate Bayesian computation [Beaumont, 2019, Sunnåker et al., 2013], among others.

Second, we are collecting HST data with well-annotated spatial domain information as the pilot data for spaDesign. This initiative aims to create an extensive atlas covering samples from diverse tissues (e.g., heart, brain, kidney, lung), species (e.g., human, mouse), and conditions (e.g., tumor, diseased, healthy). By doing so, we hope to fully exploit the full capabilities and benefits that can be offered by spaDesign.

Third, we plan to extend our framework to imaging-based and multi-sample HST experiments, each presenting unique challenges. While our current framework centers on a singular experimental design factor (total sequencing depth), imaging-based HST experiments introduce the field of view (FoV) as a critical consideration. Specifically, both the number and the size of FoV are important design factors. Moreover, as HST experiments increasingly involve the collection of multiple tissue sections from a single sample or the collection of one tissue from multiple individuals, the number of sections and/or individuals becomes another pivotal design factor.

In conclusion, our ongoing efforts to enhance the spaDesign framework are multifaceted. We hope to empower researchers with robust tools for more informed experimental design decisions, and ultimately drive the progress in the field of spatial genomics.

## Acknowledgments

We also want to express our sincere gratitude to the NIMBLE team for their invaluable assistance with the parallelization of the Poisson Gaussian process code. Additionally, we extend our thanks to the authors of the Fisher-Gaussian kernel model for permitting us to reuse their code. This work was supported by the National Human Genome Research Institute [R21 HG012482]; the National Institute of General Medical Sciences [R01 GM152585], the National Institute on Aging [U54 AG075931]; the National Institute on Drug Abuse [U01 DA045300]; the National Science Foundation [NSF1945971]; and the Pelotonia Institute for Immuno-Oncology (PIIO). The funders had no role in study design, data collection and analysis, decision to publish, or preparation of the manuscript.

