## supplementary text for "spaDesign: A Statistical Framework to Improve the Design of Sequencing-based Spatial Transcriptomics Experiments"

### Supplementary materials for "spaDesign: A Statistical Framework to Improve the Design of Sequencing-based Spatial Transcriptomics Experiments"

**Juan Xie**[1,2,3], **Hyeongseon Jeon**[4], **Won Chang**[5,], **Yeseul Jeon**[6], **Zihai Li**[3], **Qin Ma**[2,3], **and Dongjun Chung**[1,2,3,*]

[1] The Interdisciplinary PhD program in Biostatistics, The Ohio State University, Columbus, Ohio, U.S.A.

[2] Department of Biomedical Informatics, The Ohio State University, Columbus, OH, U.S.A.

[3] Pelotonia Institute for Immuno-Oncology, The James Comprehensive Cancer Center, The Ohio State University, Columbus, OH 43210, U.S.A.

[4] Department of Mathematics, University of Houston, Houston, Texas, U.S.A.

[5] Department of Statistics, Seoul National University, Seoul, Republic of Korea.

[6] Department of Statistics, Texas A&M University, College Station, Texas, U.S.A.

[*]

### 1 Impact of the number of selected spots

To illustrate the impact of the number of selected spots during conditional sampling, we checked the simulated expression for gene MBP in domain WM in the human brain data (sample 151673). Specifically, we varied the proportion of selected spots from 20% to 90% and conducted conditional sampling. The obtained simulated expression, as well as the original expression, are shown in Fig S1. We can see that when the proportion of selected spots ranges between 50% and 90%, the simulated expression closely mirrors the original pattern. However, while when the proportion drops below 50%, the simulated pattern begins to deviate from the original pattern.

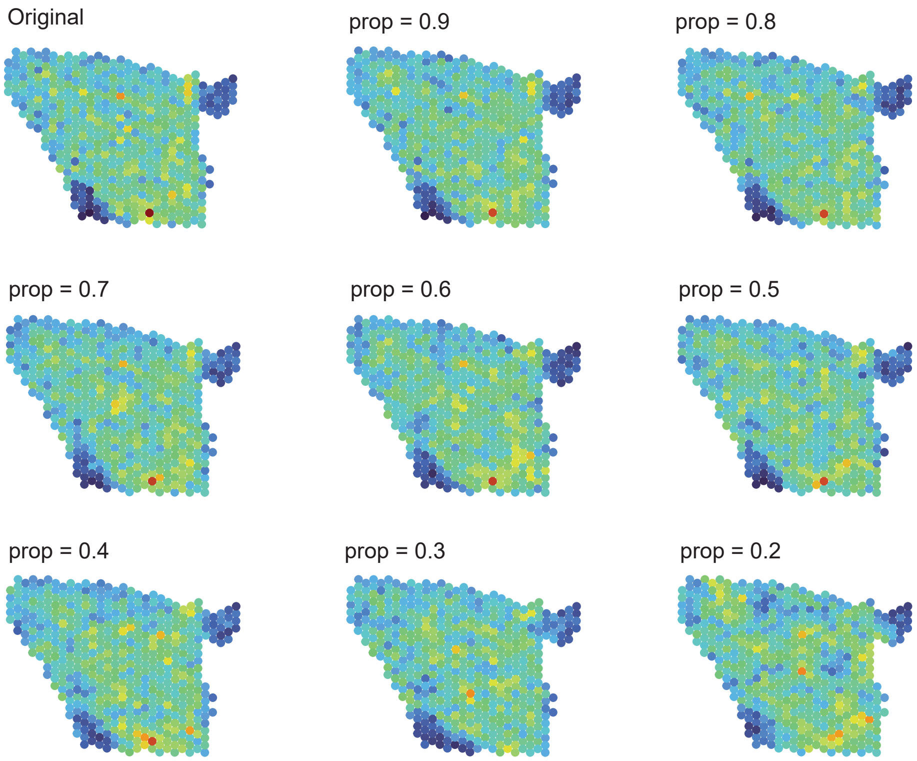


Figure S1: **Impact of the proportion of selected spots on the conditional sampling of the fitted Poisson Gaussian process.**

#### 2 Details of the spatial domain detection tools

We applied three spatial domain detection methods to detect tissue domains from the synthetic data. The three methods include BayesSpace (v1.2.1), SpaGCN (v1.2.7), and Louvain implemented in Seurat (v4.4.0), representing three major spatial domain detection methods that are commonly used for HST data. Louvain [1] is an unsupervised clustering method that detects small communities from a graph by optimizing modularity. SpaGCN [2] is a machine learning method that uses a graph convolutional network to integrate gene expression, spatial location, and histology for spatial domain identification. BayesSpace [3] is a Bayesian multivariate $t$-mixture model that utilizes hidden Markov random fields to incorporate spatial information with gene expression data to identify spatial domains. For BayesSpace and SpaGCN, we directly set the expected number of spatial domains to be the true number of domains in the pilot data. For Louvain, we first over-cluster by setting a large resolution value (res=3). Then, we cluster the cluster centers hierarchically and cut the resulting tree at the desired number of domains.

### 3 Supplementary figure for the chicken heart data

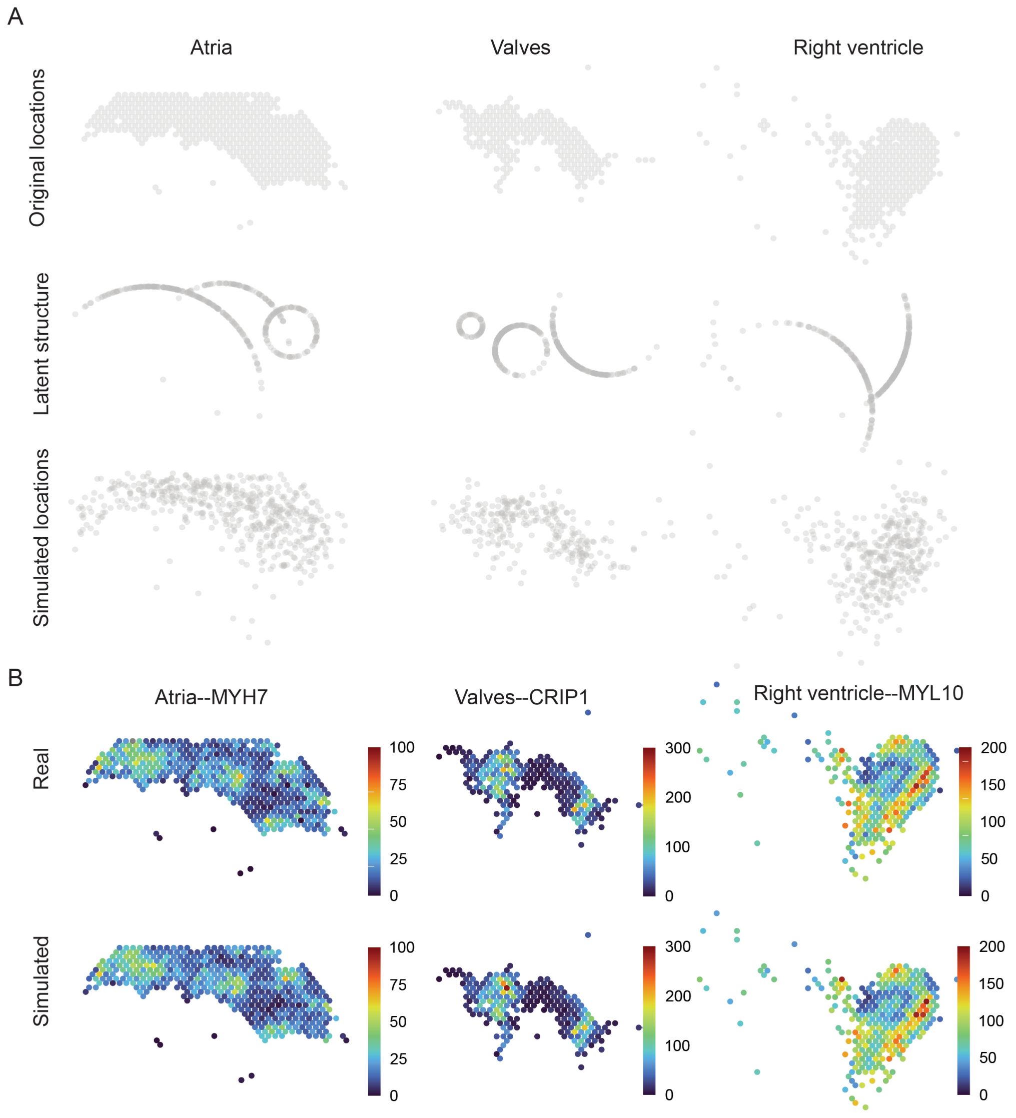


Figure S2: **spaDesign simulates data in a flexible way using the chicken heart data.** (A) The spatial distribution of points from three selected domains is displayed (the first row), along with the estimated latent structure (the second row) and the simulated locations generated using spaDesign (the third row). (B) The spatial expression patterns of three representative genes are displayed for the pilot data (the first row) and the simulated data generated by spaDesign (the second row).

### 4 Results for two additional human brain datasets

We applied spaDesign to two additional human brain datasets from the human dorsolateral prefrontal cortex (DLPFC), sample 151674 and sample 151609. Sample 151674 contains expression profiles of 33,538 genes across 3,635 spots, while sample 151609 contains expression profiles of 33,538 genes across 4,788 spots. The total sequencing depths for samples 151674 and 151609 are 21.5 million and 12.2 million reads, respectively. According to the original study, the locations are clustered into one of seven tissue domains for both samples. Based on 10X Genomics' guideline, the recommended minimum total sequencing depth would be 90.8 million read pairs for sample 151674, and 119.7 million read pairs for sample 151609. Similar to sample 151673 shown in the main text, we evaluated the impact of effect size as well as spatial pattern on the spatial domain detection performance.

#### 4.1 Impacts of effect size on the spatial domain detection performance

When the effect size is comparable to that of the pilot data, the curve made a plateau at around 3 times the original sequencing depth (Fig S3), which is approximately 64.5 million read pairs for sample 151674 and 36.7 million read pairs for sample 151609.

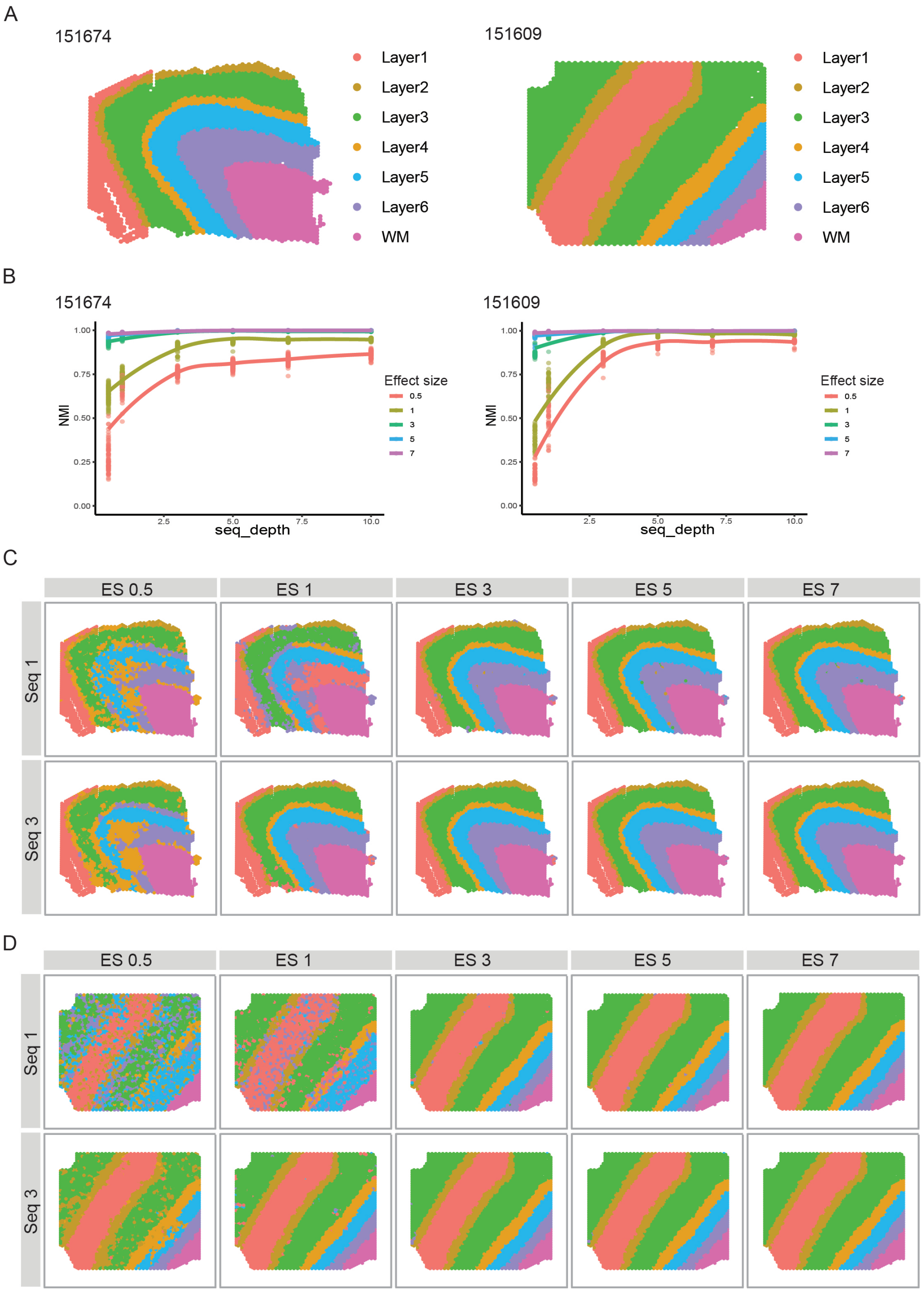


Figure S3: **Impacts of effect size on the spatial domain detection performance for the two additional human brain datasets.** (A) Manual annotations for the two additional human brain datasets. Both sample 151674 (left) and sample 151609 (right) contain seven layers, including six DLPFC layers and white matter (WM). (B) NMI curves as a function of sequencing depths ($x$ axis) under different effect sizes (lines) for human brain DLPFC sample 151674 (left) and sample 151609 (right) dataset, respectively. (C) and (D) show the example spatial domain detection results produced by SpaGCN on the simulated data under different effect sizes (columns) and sequencing depths (rows) for the two human brain datasets.

#### 4.2 Impacts of spatial pattern on the spatial domain detection performance

For the two additional human brain datasets, we observe a consistent trend in the NMI across sequencing depths (Fig S4A), similar to what we presented in the main text for sample 151673. Regardless of the proportion of disturbed genes, the NMI initially increases with sequencing depth before reaching a plateau. Moreover, there is a clear separation of the NMI curves based on the proportion of disturbed genes. Specifically, the NMI curve at a high proportion of disturbed genes is notably lower compared to those at lower proportions. As the proportion of disturbed genes decreases, the NMI curves tend to converge.

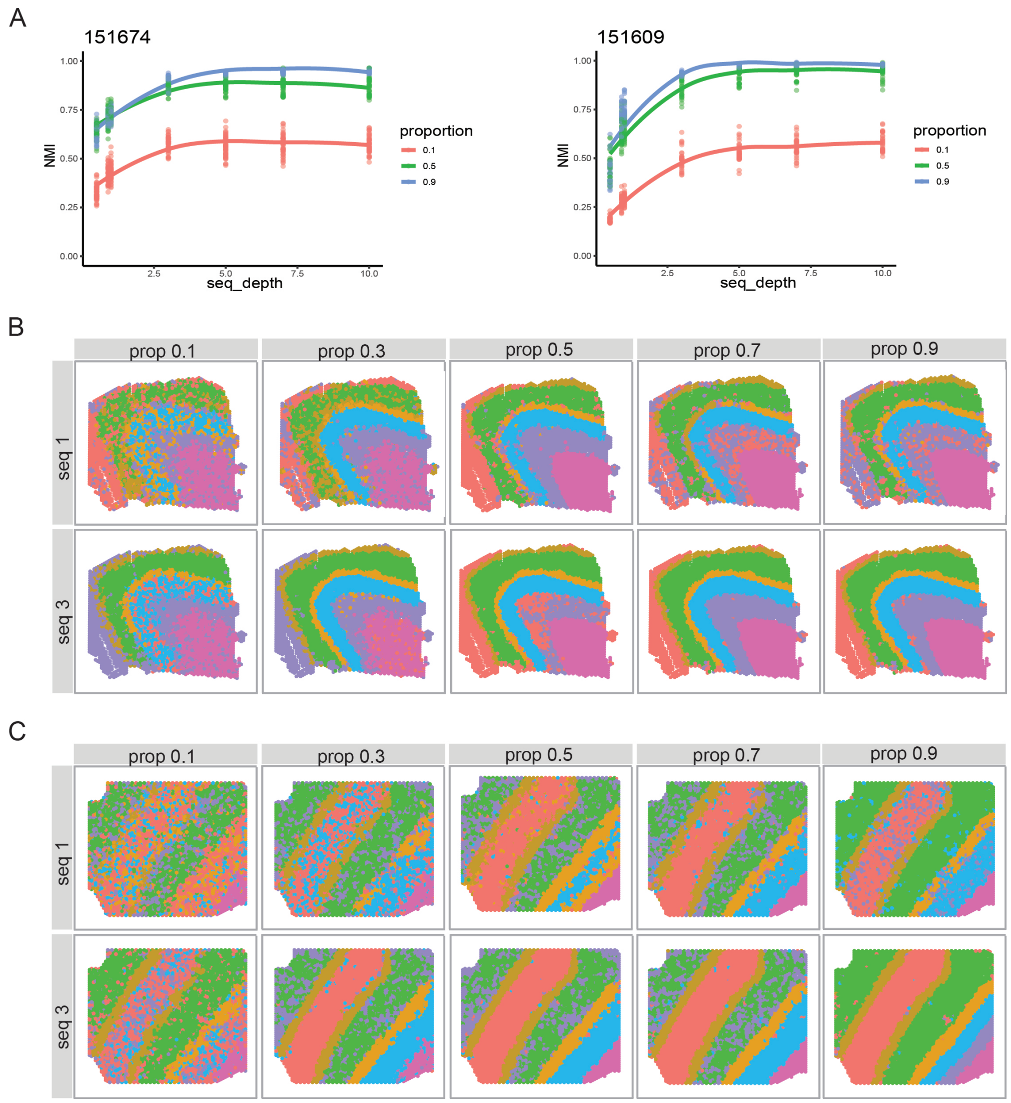


Figure S4: **Impacts of spatial expression patterns on spatial domain detection performance for the two additional human brain datasets.** (A) NMI curves as a function of sequencing depths ($x$ axis) for different proportions of disturbed genes (lines) for human brain DLPFC sample 151674 (left) and sample 151609 (right). (B) and (C) show the example spatial domain detection results produced by SpaGCN on the simulated data under different proportions of disturbed genes (columns) and sequencing depths (rows) for the two human brain DLPFC sample 151674 (B) and sample 151609 (C), respectively.

### 5 NMI curves obtained using BayesSpace and Louvain

Next, we applied two other spatial domain detection methods, BayesSpace [3] and Louvain (implemented in Seurat) [1], to the simulated datasets and obtained the NMI curves. When we compared the results between BayesSpace and Seurat under the same conditions, we found that BayesSpace generally provided more stable performance and achieved higher NMI scores than Seurat. Additionally, we observed that Seurat reaches saturation at a higher sequencing depth compared to BayesSpace. This suggests that Seurat requires a greater sequencing depth to achieve its maximum performance. This makes sense and is expected given the fact that Seurat does not utilize spatial information when detecting spatial domain. This difference may highlight the role of spatial information in spatial domain detection to potentially achieve the optimal results with a lower sequencing depth.

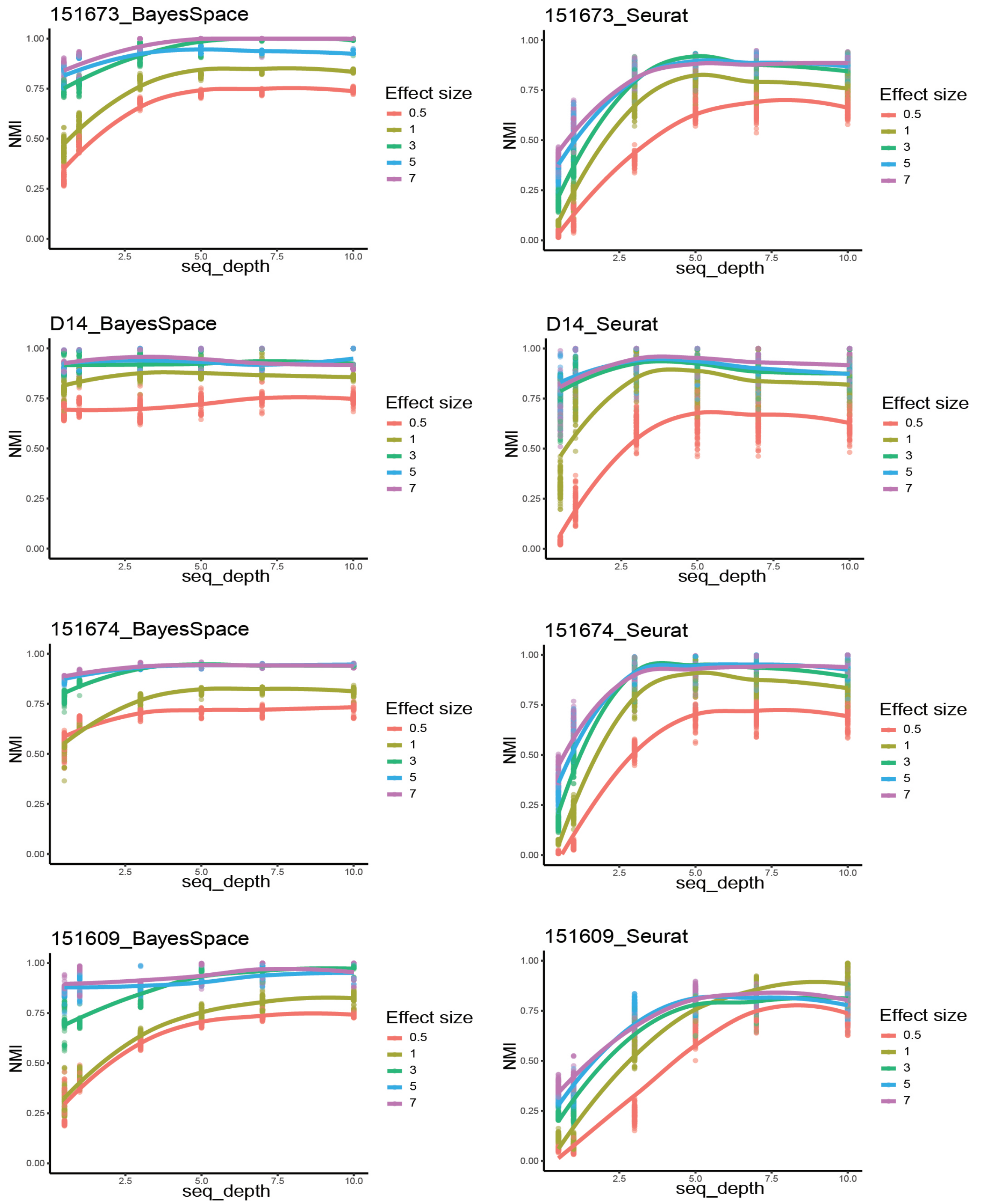


Figure S5: **Impacts of effect size on the spatial domain detection performance using BayesSpace and Louvain.** The NMI curves as a function of sequencing depths ($x$ axis) under different effect sizes (lines), where the results were obtained using BayesSpace (left) and Seurat (right).

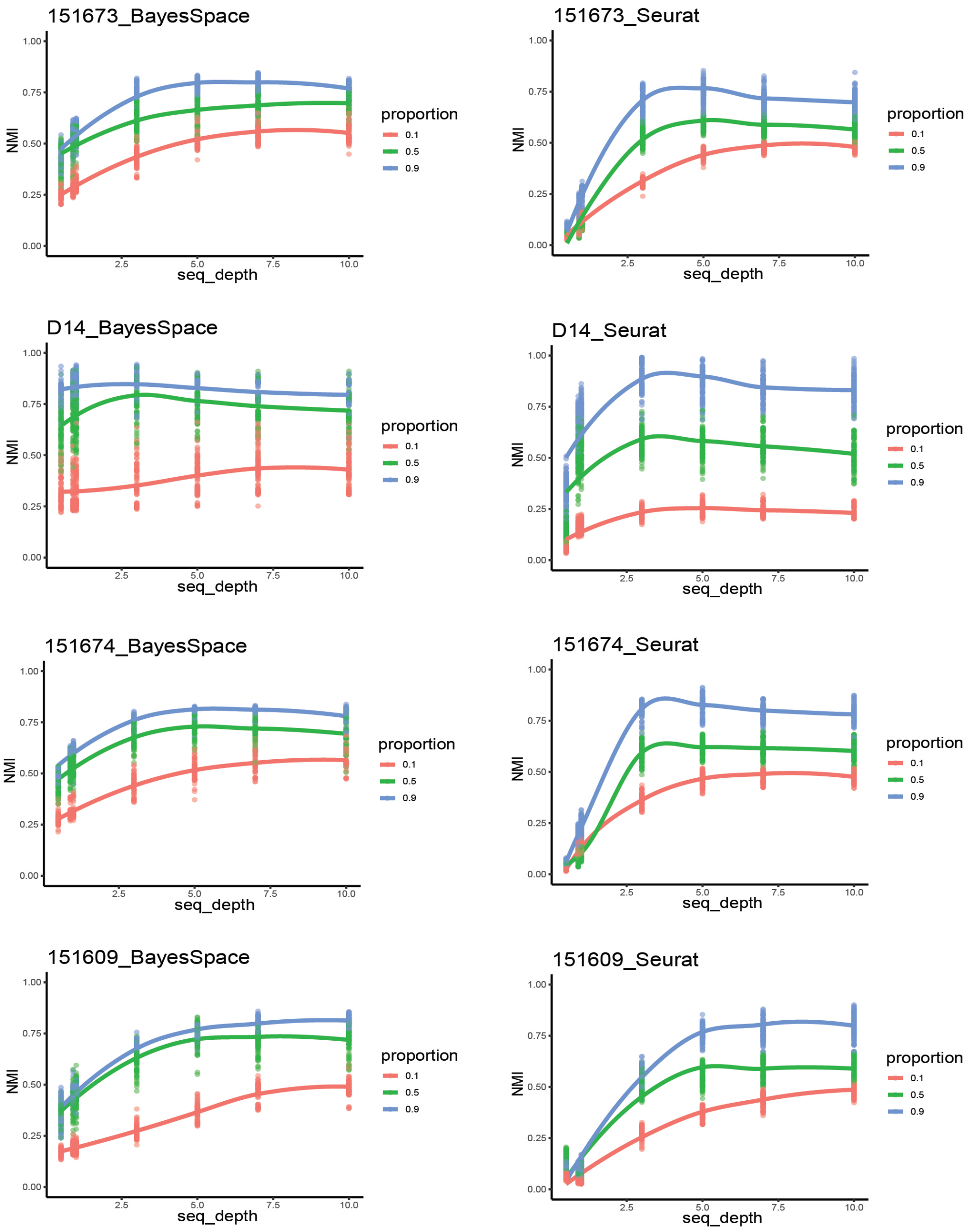


Figure S6: **Impacts of spatial expression pattern on the spatial domain detection performance using BayesSpace and Louvain.** The NMI curves as a function of sequencing depths ($x$ axis) under different proportions of disturbed genes, where the results were obtained using BayesSpace (left) and Seurat (right).
